# Sex differentiation in grayling (Salmonidae) goes through an all-male stage and is delayed in genetic males who instead grow faster

**DOI:** 10.1101/135194

**Authors:** Diane Maitre, Oliver M. Selmoni, Anshu Uppal, Lucas Marques da Cunha, Laetitia G. E. Wilkins, Julien Roux, Kenyon B. Mobley, Susanne Knörr, Marc Robinson-Rechavi, Claus Wedekind

## Abstract

Fish can be threatened by distorted sex ratios that arise during sex differentiation. It is therefore important to understand sex determination and differentiation, especially in river-dwelling fish that are often exposed to environmental factors that may interfere with sex differentiation. However, sex differentiation is not sufficiently understood in keystone taxa such as the Thymallinae, one of the three salmonid subfamilies. Here we study a wild grayling (*Thymallus thymallus*) population that suffers from distorted sex ratios. We found sex determination in the wild and in captivity to be genetic and linked to the sdY locus. We therefore studied sex-specific gene expression in embryos and early larvae that were bred and raised under different experimental conditions, and we studied gonadal morphology in five monthly samples taken after hatching. Significant sex-specific changes in gene expression (affecting about 25,000 genes) started around hatching. Gonads were still undifferentiated three weeks after hatching, but about half of the fish showed immature testes around seven weeks after hatching. Over the next few months, this phenotype was mostly replaced by the “testis-to-ovary” or “ovaries” phenotypes. The gonads of the remaining fish, i.e. approximately half of the fish in each sampling period, remained undifferentiated until six months after fertilization. Genetic sexing of the last two samples revealed that fish with undifferentiated gonads were all males, who, by that time, were on average larger than the genetic females (verified in 8-months old juveniles raised in another experiment). Only 12% of the genetic males showed testicular tissue six months after fertilization. We conclude that sex differentiation starts around hatching, goes through an all-male stage for both sexes (which represents a rare case of “undifferentiated” gonochoristic species that usually go through an all-female stage), and is delayed in males who, instead of developing their gonads, grow faster than females during these juvenile stages.

**Author contribution:** MRR and CW initiated the project. DM, OS, AU, LMC, LW, and CW sampled the adult fish, did the experimental *in vitro* fertilizations, and prepared the embryos for experimental rearing in the laboratory. All further manipulations on the embryos and the larvae were done by DM, OS, AU, LMC, and LW. The RNA-seq data were analyzed by OS, JR, and MRR, the histological analyses were done by DM, supervised by SK, and the molecular genetic sexing was performed by DM, OS, AU, and KBM. DM, OS, and CW performed the remaining statistical analyses and wrote the first version of the manuscript that was then critically revised by all other authors.

## 1. Introduction

Fishes show a great diversity of gonadal development and differentiation that can be classified into five categories: (i) gonochoristic species with individuals developing either testes or ovaries, (ii) sequential hermaphrodites that mature as males (protandrous) or (iii) as females (protogynous) and may change sex later in life, (iv) simultaneous hermaphrodites, and (v) all-female species that reproduce gynogenetically ^1, 2^. In the gonochoristic species, primordial germ cells are typically formed during embryo or early larval development and subsequently differentiate into male or female gonads under the influence of genetic mechanisms and/or endocrine, environmental, or behavioural signals ^1^. This process can be direct, as in the so-called “differentiated” gonochoristic species ^3^ where primordial germ cells develop without any detour into testicular or ovarian tissues, for example, in Arctic charr (*Salvelinus alpinus*) ^4^ and some cyprinid fishes ^5, 6^. In “undifferentiated” gonochoristic species, the typical pattern is that individuals first develop ovarian tissues that may subsequently degenerate, followed by a masculinization of the gonads that finally leads to normal testes, as in zebrafish (*Danio rerio*) and some other cyprinids ^7, 8^. Other species, so-called “secondary” gonochoristics ^9^, seem to first develop into simultaneous hermaphrodites before most individuals mature as only females or males, as in many eel populations *Anguilla* sp. ^10, 11^. We know of no example of an “undifferentiated” gonochoristic species where all individuals first develop immature testicular tissue followed by a feminization of the gonads that finally leads to normal ovaries.

Fishes also show a great diversity in sex determination systems that range from purely genetic to purely environmental, and different types of environmental sex reversals have been described in many different orders of the teleosts ^1, 12^. The fact that sex determination in fish can be very labile has several practical consequences: it can (i) be exploited in aquaculture where one-sex cultures can sometimes be more profitable ^13^, (ii) be used to control problem species, e.g. invasive species ^14, 15^, (iii) be used to boost population growth in the wild ^16^, and (iv) cause some species to be sensitive to environmental changes, especially to different types of endocrine-disrupting pollutants ^17, 18^. It is therefore important to understand the diversity of sex determination and sex differentiation among fishes.

We study a population of European grayling (*Thymallus thymallus* L.) that uses the pre-alpine lake Thun as its feeding habitat, and that spawns in spring at the outlet of this lake. A yearly monitoring program initiated in the 1940s revealed significantly distorted adult sex ratios (an excess of males) that coincide with an abrupt temperature change in Europe ^19, 20^ and that may contribute to the continuous decline of the population ^21^. Grayling belong to the salmonids that include the three subfamilies Salmoninae (e.g. Pacific and Atlantic salmon, trout, char), Thymallinae (grayling), and Coregoninae (whitefish). Salmonids are usually keystone species of their respective habitat and of considerable economic and cultural importance for local communities. Sex determination seems mostly genetic in Salmoninae and Thymalinae, where it may be driven by a master sex-determining gene ^22^. With regard to sex differentiation, much variation is observed among the Salmoninae ^1^, and little is known about the Thymalinae, including the European grayling (*Thymallus thymallus*).

Here we first verified that the *sdY* locus that Yano et al. ^22^ tested on 54 graylings from a fish farm in France can be used to predict phenotypic sex in our wild study population. We searched for genotype-phenotype mismatches (i) in wild-caught breeders, (ii) in F1 progeny that were raised to maturity in captivity, and (iii) in juvenile fish that were raised in the laboratory under warm or cold conditions. We then experimentally produced half-sib groups and raised the embryos first singly in 24-well plates, then group-wise during their larval and juvenile stages. We extracted mRNA from embryos, hatchlings, and early larvae, a time window that corresponds to the onset of sexual differentiation in rainbow trout (*Oncorhynchus mykiss*) ^23^, to detect sex-specific gene expression patterns. We also used histological techniques to describe sex differentiation in relation to genetic sex markers over a period of several months.

## 2. Methods

### 2.1 Verification of genetic sex determination and genetic sexing of larvae

For determining the sex of larvae and juveniles, genomic DNA was extracted from tissue samples (tails or fins) using the DNEasy Blood and Tissue Kit (Qiagen, Hombrechtikon, Switzerland), following manufacturer’s instructions. To verify the efficacy of the sex typing primers, fin clips were taken of in total 192 adults that were either caught from the study population ^21^ (92 males and 29 females) or F1s that had been raised in captivity until sexual maturity (20 males and 49 females). All these fish showed the typical sexual dimorphism of this species. They were stripped for their gametes, i.e. phenotypic sex could be verified without dissection. For adult tissue, DNA was extracted using a BioSprint 96 robot tissue extraction kit (Qiagen).

Polymerase chain reaction (PCR) was conducted following protocols by Yano et al. ^22^ with modifications. Briefly, each 15μL PCR contained 1.5μL 10X PCR Buffer, 210 μM dNTPs, 1.5 mM MgCl_2_, 0.3 μM each of primers *sdY E1S1* and *sdY E2AS4* (male-specific amplification ^22^), 0.075 μM each of primers *18S S* and *18S AS* ribosomal RNA (positive amplification control ^22^), 0.75 units of Taq Polymerase (Qiagen or Promega GoTaq), and 100 ng of DNA. Thermal cycling consisted of denaturing for 3 min at 95°C followed by 40 amplification cycles of 94°C/30s, 62°C/30s, and 72°C/30s, with a final extension of 10 min at 72°C. PCR products were visualized on a 1.5% agarose gel at 100V for 50min (Supplementary Figure S1).

A multiplex reaction with three microsatellite markers previously used to characterize population differentiation in grayling: *BFRO005, BFRO006* ^24^ *and Ogo2* ^25^ and with *ThySex225* was used to analyze the tissue samples of the adults. We designed a small amplicon (∼225bps) of the *sdY* locus from sequences specific to grayling ^22^ using Primer3 ^26^ (*ThySex225*: 5□ -AGCCCAGCACTCTTTTCTTATCTC-3 □; genbank probe DB accession #: Pr032825786). The 5’ end of the *ThySex225* was labelled with a fluorescent label (ATTO532) to aid viewing using capillary gel electrophoresis and to multiplex with microsatellites for high-throughput genotyping. We used the reverse primer *sdY E2AS4* of Yano et al. ^22^. Each polymerase chain reaction (PCR) was accomplished in a10μl reaction volume containing 1.5μl water, 5μl of Qiagen Hotstar Taq Mix (final concentration 0.5 units of HotStarTaq DNA Polymerase, 1XPCR buffer, 1.5mM MgCl_2_ and 200uM of each dNTP), 0.3μl of each primer (10μM), and 2μl of template DNA (1-5ng/μl). The thermal cycling profile consisted of an initial denaturation for 15 min at 95°C followed by 35 cycles of 94 °C (30 sec), 56°C reannealing temperature (90 sec) an extension phase at 72°C (60 sec), and a final extension at 72°C for 30 min. Fragment lengths were visualized on an Applied Biosystems® (ABI, Life Technologies GmbH, Darmstadt, Germany) 3730 capillary sequencer and alleles were scored using GeneMarker Version 2.6.4 software (SoftGenetics, LLC, State College, PA, USA).

### 2.2 Breeding experiments

Two sets of breeding experiments contribute to the present analyses. For the first breeding experiment, gametes were stripped from 16 wild genitors (8 males and 8 females) and used for block-wise full-factorial breeding with two males and two females per block. The embryos of these 16 half-sib groups were distributed to 24-well plates (Falcon, Becton-Dickinson), with two eggs per well that had been filled with 2 mL of chemically standardized water ^27^. After hatching, in total 532 fish were about equally distributed to eight 200L aquaria filled with filtered lake water (closed system). Fish were randomly assigned to 4 aquaria at 12°C and 4 aquaria at 18°C in separate climate chambers. At day 174 after fertilization (i.e. day 145 after peak hatching), two of the aquaria of each temperature treatment were exposed to the parasite *Tetracapsuloides bryosalmonae* in the course of another study (Uppal *et al*., unpublished manuscript). Exposure to *T. bryosalmonae* had no effect on any analysis presented here (data not shown).

Fish were fed *ad libitum*, initially on a live zooplankton and then on dry food (Skretting, Nutra Brut 3.0, 2.0, T-1.1). At days 63-71 post exposure, all fish were euthanized with an overdose of Koi Med Sleep (Ethylenglycolmonophenylether) and sexed by visual inspection of the gonads or via the sdY genotype. In total 60 individuals were both morphologically and genetically sexed to test for possible genotype-phenotype mismatches at this stage.

The second breeding experiment was performed similarly to the first breeding experiment, with the following modifications. The genitors were sampled from a captive breeding stock (F1 progeny of the wild population), and their gametes stripped and used in two full-factorial breeding blocks with 4 females crossed with 5 males each, resulting in 40 different half-sib groups. The embryos were raised singly in 2 mL wells of 24-well plates at 7°C. At 14 days post fertilization (*dpf*), embryos were exposed either to 1 ng/L 17α-ethinylestradiol (“EE2”), to *Pseudomonas fluorescens* (“PF”; 10^6^ bacterial cells per well), simultaneously to EE2 and *P. fluorescens* (“EPF”), or sham-treated (“control”) in the course of parallel studies on the evolutionary potential of the population to adapt to environmental stress during embryogenesis (Marques da Cunha, L., Maitre D. *et al*. unpublished manuscript) and on EE2-effects on gene expression at different developmental stages ^28^.

In order to induce and synchronize hatching, the temperature was raised to 10°C at 27 *dpf* and to 11.5°C the next day. At 40 *dpf*, i.e. 11 days after peak hatching, a random sample per treatment was transferred to 8 tanks filled with 200 L lake water. Two of the 8 tanks each were stocked with fish that had been either exposed to EE2, *P. fluorescens*, EE2 and *P. fluorescens*, or nothing, respectively. Once per week, 40L of water per tank was replaced with either filtered lake water (groups “controls” and “PF”) or with filtered lake water to which EE2 had been added to reach a concentration of 1ng/L (groups “EE2” and “EPF”). The larvae were fed with live *Artemia*, then live copepods and later also dry food as in the first breeding experiment. Temperature was gradually increased to 18°C towards the end of the study at 163 *dpf* (in order to simulate the increase of mean temperature in the wild). The treatment with *P. fluorescens* during embryogenesis did not show any significant effects during larval stages while treatment with EE2 delayed sex differentiation ^28^ and seems responsible for 2 cases of ovarian tissues in genetic males. EE2-treated individuals were therefore excluded from the present analyses.

Larval length was determined by digital analysis of photos taken from freshly killed fish (first breeding experiment) or from the fixed and stained sections that were used for histological analyses (second breeding experiment; see below) using ImageJ ^29^.

### 2.3 Sampling and preparations for gene expression analyses

A subset of 5 half-sib groups (1 female crossed with 5 males) from the treatment groups “controls” and “EE2” of the second breeding experiment was used to also study gene expression during early developmental stages. The first samples (12 embryos per family and treatment, i.e. 120 in total) were taken at 21 *dpf.* Embryos were immediately transferred to RNAlater (Thermo Scientific, Reinach, Switzerland) and snap frozen at -80°C. The second sampling occurred the day of hatching (31 *dpf*) with 8 larvae per family and treatment (i.e. 80 in total). The third sampling was at day 52 *dpf* (i.e. 21 days after hatching) with 5 larvae per family and treatment (40 in total). Larvae of the second and third sampling were euthanized with KoiMed (0.5 mL/L for five minutes) and then decapitated. The heads were immediately stored in RNAlater at -80°C for later analyses. We sampled heads because the neuroendocrine system is known to play a crucial role in the sexual differentiation ^1^. For example, sex differentiation of the brain is strongly dependent on the local action of estrogenic compounds ^30^.

RNA extractions were performed using the QIAgen 96 RNeasy Universal Tissue Kit (Qiagen) following the manufacturer instructions, except that the centrifugation was done at half of the protocol speed for double the amount of the time (Eppendorf 5804 R centrifuge with an A-2-DWP rotor; Eppendorf, Schönenbuch, Switzerland). In total, three distinct runs of extractions (up to 96 samples each) were performed. Samples under the same treatment, from the same family and collected at the same developmental stage were assigned to the same run of extraction. RNA was extracted from the whole egg for the first sampling date and from heads only in subsequent samples. Samples were eluted in 100 μL of RNase free water.

The RNA extraction protocol we used did not include a DNase treatment. Therefore, we amplified DNA traces inside the RNA samples to determine the sdY genotype of eggs and fry. We used two amplification protocols using the 18s gene as an internal control. The first method was used in multiplex for samples with a high amount of DNA. For the samples with low DNA content, the second PCR protocol was used in single reactions with half the amounts of the respective primers each. After genetic sexing, one female and one male per maternal half-sib group and time point was haphazardly chosen for sequencing. In one family, two females were used for the second time point because no male was found in the respective family. In another family, two males were used for the third time point each because no female could be found in the respective family.

The 60 samples designed for sequencing were checked for quality (absorbance ratios and RNA-Quality-Number, RQN) and concentration using both Nanodrop (Thermo Scientific, Reinach, Switzerland) and a Fragment Analyser (Advanced Analytical, Ankeny, USA). All samples were provided for library preparation in an equimolar concentration of 6 ng/μL in 100 μL of RNAse-free water. For each library, 50 μL were used (*i.e.* 300 ng of RNA). The libraries were prepared in one batch on a robot using the Truseq Stranded RNA protocol (Illumina, Part# 15026495 Rev. A) and multiplexing adaptors. Libraries were then sequenced on an Illumina HiSeq 2500 machine to produce 2*100 bp paired -end reads. The 60 samples were sequenced in ten lanes with six samples per lane. Both the library preparation and sequencing steps were performed at the Genomic Technologies Facility at the University of Lausanne. See Supplementary Tables S1 and S2 for further details.

### 2.4 Bioinformatics RNA-seq processing

Pairs of reads of 100 bps each were quality trimmed using fastq-mcf (ea-utils, version 1.1.2; Aronesty, 2013). Low quality reads (mean <u>Phred quality score</u> below 20) and reads containing adapter sequences were trimmed. All the retained reads were truncated and returned with a length of exactly 2*90 bps. An additional quality check using FastQC (version 0.11.2; http://www.bioinformatics.babraham.ac.uk/projects/fastqc/) showed abnormal k-mer frequencies, so we removed ten additional bps at the 5’ end of reads, returning 2*80 bps long reads. This last step was performed with a custom Python script (all scripts available at https://github.com/Oselmoni/GSD_MS), which also corrected the headers of the sequence names in order to allow compatibility with downstream tools. After these processing steps, the quality of each library was rechecked using FastQC.

Twenty-four libraries, i.e. two individuals randomly chosen across each of all 12 possible combinations of developmental stage, sex and treatment, were chosen (Table S3), and duplicated reads were removed with fastq-mcf. The 24 de-duplicated read sets were then merged and used for assembly with Trinity (version 2.0.3; Grabherr et al., 2011). To speed up the assembly, we ran Trinity with the read coverage normalization option set to 50. The assembled transcriptome statistics were provided by Trinity and custom Python scripts.

To exclude spurious transcripts from the assembly, we blasted the assembled sequences against the UniProtKB/Swiss-Prot database (version of October the 16th 2015) ^31^ using the blastx command line application (version 2.2.26)^32^. For each transcript, only the best hit was kept using a cut-off E-value set at 10^-6^. The transcripts that did not have any match were filtered out of the assembly. In total, 228,417 transcripts were kept, distributed across 52,353 genes.

We used Kallisto (version 0.42) ^33^ to pseudo-map reads to the transcriptome. Next, we summed the estimated counts of the isoforms of each gene by using a custom Python script. The estimated counts per gene were used for differential expression analysis.

Normalization factors for each library were calculated using the Trimmed Mean Method (TMM) ^34^ of the EdgeR package (version 3.12.0) ^35^. A log transformation of count-per-million (log2(cpm)) was then applied to raw expression values. Log transformed count-per-million values were used for a Principal Component Analysis (PCA) of samples ^36^. Identified outlier samples (s24, s25) were filtered out. Additionally, visual inspection of the distribution of the normalized expression values showed that one sample (s58) did not follow the trend of the other samples and was therefore filtered out.

To process the remaining 57 samples, only genes showing an expression of at least 2 cpm in at least five samples were kept (in total, 35,348 genes). Two more PCAs were performed: one to investigate the role of experimental factors (developmental stage, sex, sibgroup, sequencing lane, batch of RNA extraction) in the expression measures of control individuals, and a second one applied to the whole expression matrix (i.e. with both control and EE2 treated individuals). This last PCA allowed us to select the most important factors to include in our model. We considered developmental stage, sex and treatment as a combined variable (with twelve possible levels) and the sib-group as an independent variable. This model included sufficient replication for each comparison: including sib-group inside the “combined block” would not have allowed comparisons between the different levels of this block. For this same reason, technical factors (sequencing lane and batch of RNA purification) were not included in the model.

The differential gene expression analysis was performed using the Bioconductor limma-voom package (version 3.26.3) ^37, 38^. We used the *voomWithQualityWeight*, which calculated sample quality weights in addition to the observational weights of limma-voom ^39^. We used a modified version of the *voomWithQualityWeight* function performing the cpm transformation using the cpm function from the edgeR package, resulting in a reduced range of minimal expression values across samples (since the minimal cpm values for each sample depend on the size of the library). *voomWithQualityWeight* was run with a cyclic-loess normalization step and the distribution of the expression values was manually checked (Figure S2) before proceeding to the differential gene expression analysis.

A linear model was fit for each gene, and coefficients and standard errors were computed for all contrasts of interest using limma. At each developmental stage, we obtained p-values for each gene for the comparison between control males and control females. Q-values ^40^ were obtained from the vector of p-values. A threshold of q=0.15 was applied to select differentially expressed genes, i.e. a 15% false discovery rate.

In order to have a biologically meaningful annotation of Gene Ontology (GO; Ashburner *et al.*, 2000) terms, we performed a similarity search via blastx command line application (version 2.2.26; National Center for Biotechnology Information, 2008) of the remaining transcripts against a customized database. This customized database contained all the protein sequences from Uniprot (version of December the 15th 2015; Boeackmann *et al.*, 2003) associated to Zebrafish. According to the transcript identifiers of the Trinity assembly, we were able to identify isoforms from the same gene and to keep all the associated Uniprot entries. This information was then processed by a custom Python script: for each gene, we retrieved a list of all the GO terms associated to all blast hits of the respective isoforms. Our annotation concerned only the Biological Function domain of the GO term classifications (Ashburner *et al.*, 2000). The enrichment analysis of GO terms was performed using the goseq package (version 1.22.0; Young *et al.*, 2010) of the R Bioconductor (version 1.20.1; Huber *et al.*, 2015). The list of genes differentially expressed in a contrast were checked for enrichment of GO terms using a Wallenius hypergeometric distribution inspired method (Young *et al.*, 2010). A key feature of goseq is that it takes into account the length of the gene to calculate the enrichment scores (Young *et al.*, 2010). Length of genes was measured by a custom Python script. When a gene showed multiple isoform, the median isoform length was used as length of the gene. For each GO term, goseq returns a p-value associated to the score of the enrichment test. These p-values were used to filter and rank GO terms significantly overrepresented in the gene list of interest. The results of the enrichment analysis where then visualized with REVIGO (Supek *et al.*, 2011). For each GO terms enrichment analysis, the REVIGO visualization concerned only the GO terms with a p<0.05. To assure output readability, if the number of terms having a p<0.05 was above 150, only the 150 GO terms with the lowest p-value were kept for the REVIGO visualization.

### 2.5 Gonadal development

In total 256 fish were randomly sampled 51, 79, 107, 135, and 159-163 *dpf*, i.e. about monthly over the course of 5 months. These fish were euthanized with an overdose of KoiMed Sleep, the heads stored in RNAlater at -80°C for further analyses, and the rest of the bodies fixated for two weeks in Davidson solution (AppliChem product No. A3200) for histology. They were transferred to embedding cassettes and dehydrated for 48h using a Leica TP1020 tissue processor (Leica, Tempe, USA). Dehydrated tissues were flowed in hot paraffin wax (Histoplast P, Serva, Heidelberg, Germany) using a paraffin dispenser embedding (Leica EG1150H, Leica, Tempe, USA) and paraffin wax was finally cooled to obtain a solid paraffin block. Sections were cut at 4 μm ventrally, from anterior to posterior, floated in a water-bath, and collected onto glass slides. Sections were stained with standard Mayer’s haematoxylin and eosin staining (HE-stain ^41^) and cover slipped to be conserved. Fish sections were analysed by light microscopy using a Leitz Aristoplan microscope (Leitz, Wetzlar, FRG) and analysed with an associated digital camera (Color View, Soft Imaging Systems, Münster, FRG) supported by “Analysis software” (Soft Imaging Systems, Münster, FRG).

## 3. Results

### 3.1 Verification of genetic sex determination

Using the microsatellite multiplex with the *Thysex225* locus, we assigned sex to 121 wild-caught adults and 71 adults from the captive breeding stock. Males have a single peak at 228 bps at this locus whereas females only showed signals for the microsatellite loci. There was perfect alignment of sex assigned in the field based on morphology and on the presence of eggs and sperm with the multiplex protocol, i.e. there was no ambiguous assignment of sex based on this procedure.

Genetic sexing of 237-245 *dpf* old juveniles (first breeding experiment) based on the multiplex PCR using *sdY* primers, and 18S primers as a positive amplification control, also provided a perfect alignment between gonad morphology and the multiplex protocol with 60 individuals.

### 3.2 Gene expression

RNA quality check, sexing PCR results, RNA sequencing reads quality check, transcriptome assembly statistics and data preparation for differential gene expression analysis are given in the Supplementary Material. Figure 1 shows the results of a principal component analysis (PCA) on the normalized expression levels matrix of the 28 samples. Gene expression clustered first by developmental stage, and second by maternal half-sibgroups within each developmental stage (Fig. 1). Inside each developmental stage, sex appeared to best explain the variance in gene expression in the first two principal components in hatchlings (Fig. 1a). This is confirmed in Table 1 that reports the number of genes differentially expressed between males and females at each developmental stages. While only 15 genes were differentially expressed at the embryo stage (Fig. S3), a strong increase of differently expressed genes could be observed at the day of hatching (72% of total genes, Fig. S4). The sex difference remained important at the first feeding stage (3% of total genes, Fig. S5). The GO enrichment analysis of the genes differentially expressed between males and females are shown in the supplementary figures S6-S9.

**Table 1.** Number of genes differentially expressed (q<0.15) between males and females at three developmental stages.

| Developmental stage | Number of genes |
| --- | --- |
| Embryo | 15 |
| Hatchling | 25,372 |
| First feeding | 1,110 |

**Figure 1.**
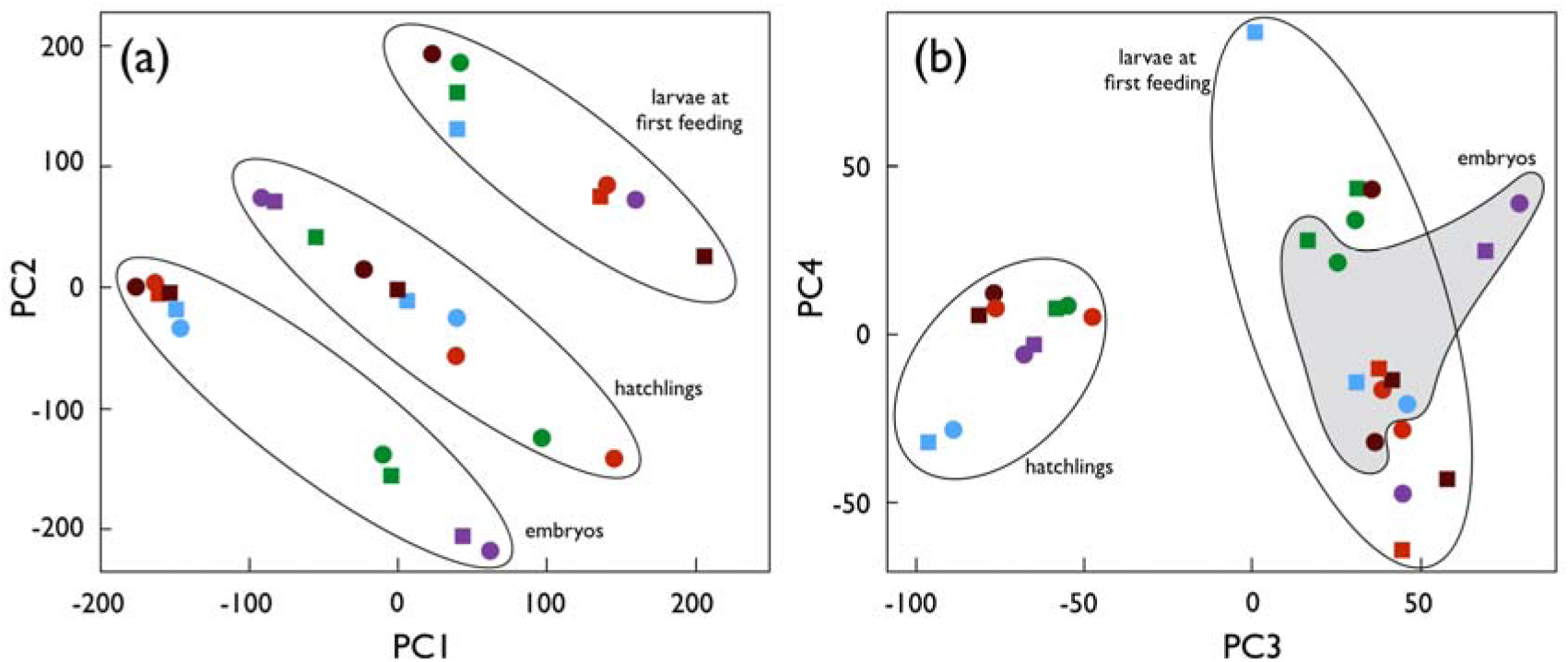
Principal component analysis of the gene expression matrix. (a) The first two principal components, explaining 35.0% and 34.7% of the observed variance, respectively, and (b) the next two principle components, explaining 9.5% and 3.3% of the variance, respectively. Round symbols represent females, squared symbols males, the colours represent the 5 maternal half-sib families. Ellipses and the grey area in panel b emphasize the clustering by developmental stage.

### 3.3 Gonadal development

Figure 2 gives an overview of the sex differentiation over the 5 sampling periods of the second breeding experiment (Fig. 2). The figure also gives the frequency of genetic males and females in the first and the second breeding experiment. All gonads were undifferentiated at the first sampling period (51 *dpf*). The rate of undifferentiated gonads dropped to 30.4% at the second sampling period (79 *dpf*). The other fish first showed early testicular tissues while there were no signs of ovarian tissue. From the second to the third sampling day (107 *dpf*), the rate of fish with testes only dropped from 69.6% to 20.8%, while 45.8% of the fish showed testis-to-ovary or ovaries only. From then until the fifth sampling, the rate of fish with undifferentiated gonads did not decline further, and there was no clear change in the rates of fish with different types of gonadal tissues varied. All fish that remained undifferentiated at the fourth and fifth sampling period turned out to have the male genotype, while all except three of the remaining fish showed the females genotype (Table 2). See Figure 3 for examples of the various phenotypes.

**Table 2.** Gonads at sampling periods 135 *dpf* and 159-163 *dpf* (both sampling periods pooled) versus result of genetic sexing.

| Gonads | Genetic sexing |  |
| --- | --- | --- |
|  | Males | Females |
| Undifferentiated | 22 | 0 |
| Testicular tissue only | 3 | 3 |
| Testis-to-ovaries | 0 | 14 |
| Ovaries | 0 | 5 |

**Fig. 2.**
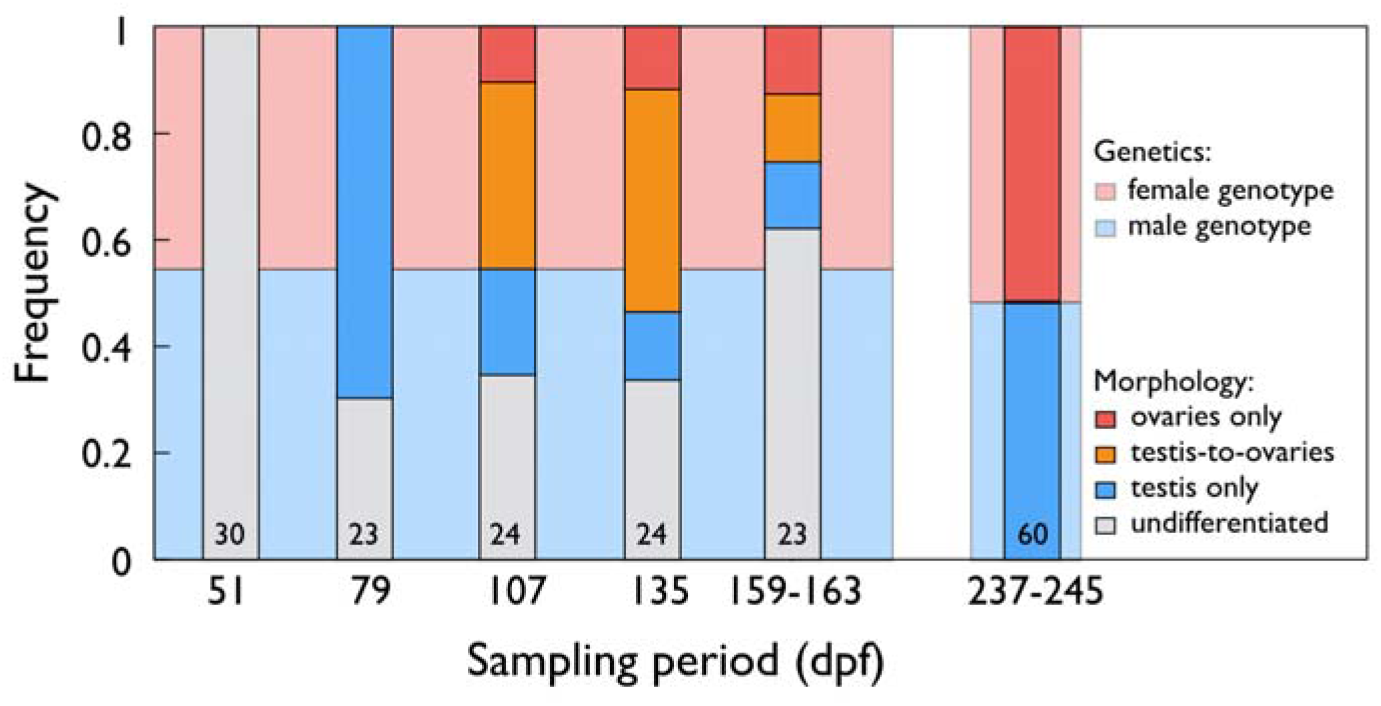
Sex differentiation in grayling. Frequencies of fish with undifferentiated gonads (grey bars), gonads with testicular tissue only (blue bars), the testis-to-ovary phenotype (orange bars), and ovaries only (red bars) for the second experiment (sampling periods between 51-163 *dpf*) when phenotypes were determined by histology, and in the first experiment (sampling period at 237-245 *dpf*) when phenotypes were determined by morphology. The numbers in the boxes give the total sample sizes for the second experiment, and the number of fish that were both phenotypically and genetically sexed for the first experiment. The blue and red background colors indicate the overall frequencies of genetic males and females, respectively, for each of the two experiments. Phenotype and genotype matched perfectly for the first experiment (sampling period 237-245 *dpf*). At the last two sampling periods of the second experiment (135 *dpf* and 159-163 *dpf*), all fish with undifferentiated gonads had the male genotype, 3 of 6 individuals with testes had the male genotype, all other individuals had the female phenotype. See table 2 for the match between phenotype and genotype at sampling periods after 135 *dpf* and Fig. 3 for examples of the various developmental stages.

**Fig. 3.**
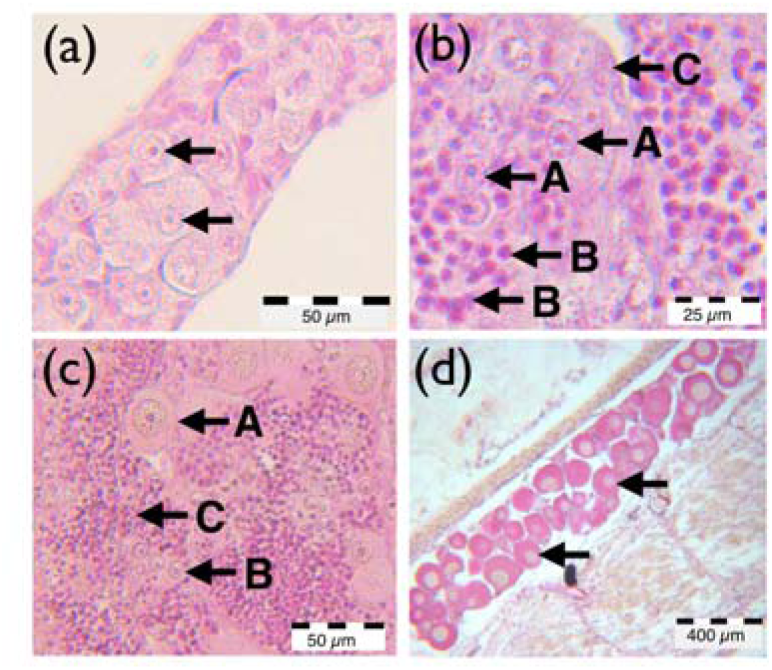
Representative examples of (a) undifferentiated gonads at fifth sampling period (arrows mark oogonia), (b) immature testis at second sampling period (A = spermatogonia, B = spermatocytes, C = Sertoli cells), (c) testis-to-ovary at fourth sampling period (A = perinucleolar oocytes, B = spermatogonia, C = spermatocytes), and (d) ovary at fifth sampling period (perinuclear oocytes). Samples were fixed in Davidsons, embedded in paraffin and HE-stain.

### 3.4 Sex difference in growth

In the first breeding experiment (sampling between days 237 and 245 *dpf*, i.e. 208-216 post hatching peak), males were larger than females (on average 56.5 mm vs 54.9 mm; 95% CI = ±1.8 each, N_total_ = 436) while rearing temperature had no significant effect on size (effects of sex: t = 2.4, p = 0.016; of rearing temperature: t = -0.8, p = 0.45). Males were also heavier than females (t = 1.9, p = 0.05). These findings were confirmed in the second experiment were genetic males were significantly larger than genetic females at the last two sampling periods: male trunks were on average 40.6 mm (95% CI = ±5.3) long, while female trunks were on average 35.3 mm (95% CI = ±5.8) (t = 2.7, p = 0.008).

## 4. Discussion

It is important to understand sex determination and sex differentiation especially in fish that play key roles in their respective environment ^42^, that are, or could potentially be, important in aquaculture ^13^, or that already suffer from distorted sex ratios as repeatedly observed in river-dwelling salmonids ^43^. All of this is true for the European grayling, and especially for the population we study ^21^. So far, little was known about sex determination and differentiation in this species ^22, 44^. As a rule, sex determination is generally more diverse and also more labile in ray-finned fishes and amphibians than it is in birds and mammals ^45, 46^. Sometimes, variation in sex determination can even be found within a species or a genus ^47, 48^. Therefore, even though Yano *et al*. ^22^ found genetic sex determination in 54 grayling sampled from a fish farm in France, it seems necessary to verify their finding in a geographically distinct wild population. We therefore first tested whether sex determination in a wild and in a related captive population is indeed genetic. Our multiplex protocols resulted in perfect alignment of sex phenotype and genotype in 192 adults of wild and captive origin, and in 60 juveniles that had been raised at cold and warm temperatures for 8 months. This demonstrates that sex determination has a strong genetic basis in our study population. It also supports Yano *et al*. 22’s hypothesis that the *sdY* locus is conserved among two of the three subfamilies of the Salmonidae, namely the Salmoninae and the Thymallinae, and it supports the conclusion of Pompini *et al*. ^44^ that distorted sex ratios in grayling are not due to temperature-induced sex reversal under ecologically relevant conditions. Because of the clear pattern we found, we were able to use the *sdY* locus to study sex-specific gene expression at embryonic stages, and sex-specific gonadal development at early juvenile stages.

Sex differentiation is expected to be largely controlled by steroids and gonadotropins, produced mainly by the brain in early life, and later also by gonads ^1^. We found that only a few genes show sex-specific expression in late embryogenesis. However, we cannot exclude the possibility that, by sampling whole embryos instead of heads only, we diluted, and therefore potentially missed, some sex-specific gene expression in the brain. At the time of hatching and when sampling heads only, we found that a very high number of genes showed sex-specific expression. The number of differentially expressed genes dropped towards first feeding but was still high around that developmental stage. Our findings support Baroiller *et al*. ^49^ who concluded for another salmonid, the coho salmon (*O. kisutch*), that the maximum sensitivity to an exogenous estrogenic treatment was around hatching time.

We analyzed gene expression in five different paternal half-sib groups that had been experimentally bred and raised to differ only in their paternal contribution, i.e. they differed only genetically (they all shared the same mother and were raised in the same environment). As we sampled only one female and five male breeders, a reliable quantification of the additive genetic effects on gene expression is not yet possible. However, our principle component analysis suggests that family effects are important. This suggests that there is significant heritability in gene expression around hatching. It remains to be tested whether rapid evolution in response to anthropogenic changes of the environment is therefore possible.

The first microscopically detectable characteristic of gonadal differentiation is the migration of the primordial germ cell ^50^. At the time of hatching, the number of primordial germ cells in salmonid gonads is small ^50^. When we analyzed gonadal tissue three weeks after hatching, all gonads were still undifferentiated. Four weeks later, about half of the juveniles showed testicular tissues while the remaining fish showed undifferentiated gonads. The rate of undifferentiated fish remained approximately constant over the next three sampling periods up to about the 19^th^ week after hatching, i.e. the rate of differentiated fish remained about constant, too. Surprisingly, however, the male phenotype was increasingly replaced by female phenotypes among the differentiated fish over the 14 weeks that were covered by the second to the fifth sampling periods.

Within the Salmoninae, differentiated gonochorism where primordial germ cells develop directly into testis or ovarian tissues has been described in, for example, Arctic charr (*Salvelinus alpinus*) ^4^, brook charr (*Salvelinus fontinalis*) ^50^, brown trout (*Salmo trutta*) ^51^, rainbow trout (*O. mykiss*) ^52^, and coho salmon (*O. kisutch)* ^53^. The European grayling as representative of the Thymallinae instead shows a rare form of undifferentiated gonochorism, since undifferentiated gonochoristic species usually go through an all-female stage before they differentiate into testis and ovaries ^1^. We conclude from our observations that the European grayling goes through an all-male stage before developing mature testis and ovaries.

By the end of the observational period, all fish that still showed undifferentiated gonads had the male genotype and had also grown faster, i.e. genetic males differentiated later and reached larger body lengths and body weights than genetic females. Delayed male gonad development was also observed in sea trout (*Salmo trutta* morpha *trutta* L.), but sex-specific growth was not observed during the early juvenile stages of this species ^51^. As we experimentally raised fish at warm and cold temperatures in our first experiment, we could also test for an interaction between sex-specific growth and water temperature. Such an interaction would potentially help explain the sex-specific juvenile mortality that seems to contribute to the observed correlation between increased water temperatures and population sex ratio ^21, 44^. However, we could not find any such effects of temperature on sex ratio under our laboratory conditions.

Among the 6 individuals with testis at the last two sampling times, three individuals were genetic females and would possibly have developed ovaries later. All fish with female gonads (testis-to-ovaries and ovaries) were genetic females. No genetic male showed female gonadal tissue. As mentioned above, our study population suffers from male-biased population sex ratios that may contribute to the continuous decline of the population ^21^. Similar observations have been made in other salmonid populations ^54^. Pollution of aquatic systems by hormone-active substances is probably ubiquitous wherever humans live (e.g. pollution by EE2, the synthetic component of the contraceptive pill), and there is no reason to assume that lake Thun and the spawning place that is located within the city of Thun are an exception. Various cities, villages, and different types of industries discharge their sewage into the lake or into rivers that feed the lake, normally after treatment in sewage plants that can, however, not be fully effective. Moreover, some of the lake’s whitefish populations (*Coregonus* sp.) have been observed to suffer from increased prevalence of gonadal misdevelopments ^55, 56^ that may be due to anthropogenic disturbances of the ecosystem. However, we found no phenotype-genotype mismatch in any of the adult fish that had been sampled in the wild and in the F1 breeding stock. This suggests that environmental sex reversal and its possible effects on the next generation ^17^ do not, by themselves, explain the distorted population sex ratios. It remains to be tested whether pollution by hormone-active substances and other anthropogenic changes of the environment lead to sex-specific mortality during early developmental stages, possibly because of sex-specific life histories, especially the sex-specific growth patterns and timing of gonadal differentiation that we observed here.

## 5. Acknowledgements

We thank B. Bracher, U. Gutmann, C. Küng, R. Mani, M. Schmid, and T. Vuille from the Fishery Inspectorate Bern for permissions and provision of fish for tissue sampling and experimental fertilizations, the Pachtverein Thun for catching the wild fish, the Saint-Sulpice pump station for zooplankton supplies, T. Braunbeck for access to the histology laboratory, and C. Berney, T. Bösch, J. Buser, I. Castro, E. Clark, B. des Monstiers, M. dos Santos, J. Kast, C. Luca, Y. Marendaz, D. Nusbaumer, E. Pereira Alvarez, M. Pompini, A.-L. Roulin, V. Sentchilo, R. Sermier, E. Vermeirssen, B. von Siebenthal, D. Zeugin, and the staff at Ecogenics GmbH, Switzerland, for assistance in the field or the laboratory. This project was authorized by the veterinary authorities of the cantons of Bern (BE118/14) and Vaud (VD2710; VD2956) and financially supported by the Swiss Federal Office for the Environment and the Swiss National Science Foundation (31003A_159579).

